# Multi-omics highlights challenges in assessing the composition and performance of microbial consortia for commercial applications

**DOI:** 10.1101/2025.02.26.640401

**Authors:** Derek D. N. Smith, Renuka M. Subasinghe, Caitlin Kehoe, Daniel S. Grégoire

**Affiliations:** Environment and Climate Change Canada, Ottawa, ON, Canada; Carleton University, Department of Chemistry, Ottawa, ON, Canada

## Abstract

The use of commercial microbial consortia in bioremediation is a promising method for addressing environmental pollution. These consortia are comprised of complex communities that include unculturable species that make it challenging to optimize consortia performance and carry out risk assessments for regulatory purposes. In this study, we provide a framework for using multi-omics to monitor the composition and performance of an aerobic ammonia oxidizing consortium in development for wastewater treatment. Long-read sequencing showed the consortium was dominated by an unclassified *Nitrosospira* species with the capacity for ammonia oxidation with many lower abundance taxa displaying the potential for denitrification. Considerable shifts in community composition and nitrogen cycling occurred when the consortium was grown along a redox gradient representative of wastewater for eight weeks. All aerobic and anaerobic cultures produced ammonia during the first four weeks and only aerobic cultures decreased ammonia concentrations after that time. Shotgun metagenomic sequencing showed the key ammonia oxidizing *Nitrosospira* sp. population decreased in abundance in aerobic cultures yet remained dominant in anaerobic cultures. Shotgun metatranscriptomic sequencing revealed that aerobic cultures decreased ammonia oxidation activity during the incubation and taxa that precluded detection in the starting material likely contributed to denitrification in anaerobic cultures. Metatranscriptomics showed that deamination of amino acids was an unexpected contributor to ammonia production that could negatively affect consortium performance. This study highlights how multi-omics provides insights that can be used to optimize performance and carry out risks assessments for consortia applied in different environmental settings.

**Importance:** The use of microbial consortia from diverse environments is gaining traction in terms of advancing a more sustainable bioeconomy. Optimizing consortia for different applications and ensuring they are compliant with environmental regulations is difficult because current practices rely on growing microbes with unknown physiological requirements. In this study we apply leading-edge sequencing approaches to develop a framework that addresses these challenges using a consortium in development for ammonia removal from wastewater. We demonstrate that long-read DNA sequencing provides complete genome assemblies and functional insights into key populations involved in ammonia removal that are poorly represented in taxonomic databases. We show that coupling DNA to RNA sequencing provides valuable information on changes in composition and metabolic activity that can occur under environmentally representative conditions for wastewater. Ultimately, our approach serves as an example of cutting-edge genomics applications for stakeholders to consider in developing microbial consortia for safe and effective use across diverse applications.

## Introduction

Microbial communities consist of multiple microorganisms that rely on metabolic handoffs and symbiotic relationships to live in diverse habitats. These relationships can be leveraged towards societal benefits such as improving agricultural yields through the use of natural products, protecting human health with probiotics, and using microbial detoxification pathways to remove contaminants from aquatic and terrestrial ecosystems (1–6). Much of our understanding on using microbial pathways in those contexts stems from studies with well-characterized isolates and defined mixtures of strains. Although these studies are integral to providing mechanistic insights to support optimization efforts, cultivating stable consortia from environmental sources provides several advantages for stakeholders invested in scaling up sustainable biotechnology that can work in less controlled environments.

The cultivation of stable microbial consortia can preserve the complex metabolic interactions that occur *in situ*, which can be integral to maintaining the activity of pathways for environmental applications (*e.g.,* dechlorinating microbial consortia that rely on acetogens and methanogens to support reductive dechlorination) (7). Consortia cultivation allows for more environmentally relevant mechanistic studies by supporting stronger positive selection for key taxa with desired phenotypes under environmentally-relevant conditions (8). Recreating these complex assemblages using defined mixtures of pure strains relies on all key members being cultivable, which poses a significant challenge because most microbes from environmental systems remain uncultivated (9, 10).

Consortia can be established through top-down approaches using environmental microbial communities as source material and continuously growing them under a pre-determined set of relevant conditions. Consortia can also be rationally defined using bottom-up approaches. Bottom up approaches can rely on synthetic biology tools to introduce metabolic functions of interest including through genetically tractable and modified strains and by controlling competition between different members within mixed communities (11). Both approaches present unique challenges that regulatory agencies must consider when assessing the risk of microbial products.

Synthetic biology can optimize desirable metabolic functions in scalable platforms but raises concerns about how genetically modified organisms can destabilize ecosystem services through competitive interactions and the introduction of mobile genetic elements (12, 13). Microbial consortia developed with top-down approaches raise similar concerns about the stability of commercially available mixtures maintained over time. Manufacturers must ensure stable composition of consortia to preserve desired metabolic activity across environmental conditions that can deviate from optimized growth conditions (14, 15). Regulators must also consider the stability of microbial consortia and their genes to evaluate the risks of consortia negatively affecting environmental and human health (12, 13, 16).

The main regulatory bodies tasked with assessing and approving commercial microbial remediation products in North America are jointly Environment and Climate Change Canada (ECCC) and Health Canada (HC) for Canada and the United States Environmental Protection Agency (US EPA). The Canadian Environmental Protection Act, 1999 (CEPA) is the main statutory instrument through which microbial biotechnology products are regulated in Canada wherein microbial products are designated as notifiable substances containing living organisms (https://www.canada.ca/en/environment-climate-change/services/managing-pollution/evaluating-new-substances/biotechnology-living-organisms.html). Pure cultures of microorganisms and consortia are subject to regulatory approval and notification with ECCC/HC to be placed on the Domestic Substance List (DSL) with species level identification of pure strains and community members. Consortia in this legal context only applies to natural combinations of organisms whereas synthetic communities require each constituent to be classified as a substance and be subject to regulatory approval, if not on the DSL. In comparison to ECCC/HC, the US EPA focuses on regulating “new” organisms that have undergone genetic modifications to introduce novel functions in their regulatory framework via the Toxic Substances Control Act with a large list of organisms (https://www.epa.gov/regulation-biotechnology-under-tsca-and-fifra/tsca-biotechnology-regulations).

Both processes rely on strain-level information obtained through cultivation work, which limits their ability to assess the risks tied to consortia that cannot be separated into cultivable isolates. Currently, Canada’s DSL has five consortia used for groundwater bioremediation that have completed the regulatory process in comparison to 73 microbial strains added to the DSL via a new substance notification or included based on historical usage within Canada (17, 18) (<u>Substances Search - Canada.ca</u> Accessed 2024/11/08). Addressing this policy gap is critical to ensure regulators are equipped to assess the risks for microbial-based products for consumer and industrial applications gaining more interest worldwide (5, 19, 20).

Cultivation-independent methods using short read 16S amplicon sequencing can help address these gaps, but these methods are limited because they lack the resolution required for strain-level characterization and do not provide functional information to assess consortia performance (21, 22). Emerging long read metagenomic and metatranscriptomic sequencing are promising methods that can address the limitations of amplicon sequencing. Metagenomics can provide valuable insights into community metabolic potential and further validate taxonomy. These data can be leveraged alongside metatranscriptomics to identify which consortia members are metabolically active under a range of environmentally relevant conditions.

Our objective in this study is to demonstrate how multi-omics methods can be used to characterize the composition and performance of microbial consortia to aid research and development and regulatory assessments. We used a model ammonia-oxidizing consortium being developed with a top-down approach for wastewater treatment plant (WWTP) settings. This consortium was chosen because WWTPs have served as models for community characterization using multi-omics including multiple studies that use well-defined genetic targets to characterize ammonia (NH_3_) and more broadly nitrogen cycling (23, 24).

In this work, we assess community composition and performance primarily through the lens of the key guilds required for NH_3_ removal (*i.e.,* nitrification) in WWTP: Ammonia oxidizing-bacteria (AOB) and nitrite-oxidizing bacteria (NOB) that rely on metabolic handoffs to carry out nitrification (25). We compare how different DNA extractions techniques affect the taxonomic composition and functional characterization of this nitrifying consortium in development. We test how growing this consortium along a redox gradient representative of WWTPs affects nitrogen cycling and link changes in performance to shifts in community composition using short-read metagenomic sequencing. Additionally, we explore how metatranscriptomics can be integrated into these experiments to assess consortium performance and support risk assessment. We conclude by providing examples of how these types of multi-omics frameworks could be adopted by regulators to support a paradigm shift away from relying on isolate cultivation.

## Results and Discussion

### Compositional and functional characterization of the starting consortium material

We identified 64 high quality metagenome-assembled genomes (MAG) from a single lot of the nitrifying consortium using a co-assembly approach that combined the long-reads from two DNA extraction kits used on fresh and frozen aliquots. All MAGs were classified to the Kingdom Bacteria with 13 unique phyla detected (sorted alphabetically): *Acidobacteriota, Actinomycetota, Armatimonadota, Bacteroidota, Bdellovibrionota, Chloroflexota, Deinococcota, Desulfobacterota_D, Eremiobacterota, Gemmatimonadota, Planctomycetota, Pseudomonadota,* and *Verrucomicrobiota*. A total of 42 unique families were identified, which included 16 unnamed families from the GTDB database (see sheet #1 in **Supplementary Data 1**). We also identified 46 unique genera, including 18 from unnamed genera (see sheet #1 **Supplementary Data 1**). Despite 60 of 64 MAGs recovered being 90% complete or higher, 10 MAGs were classified to unidentified genera and 59 of the 64 MAGs could not be classified to the species level (see sheet #1 in **Supplementary Data 1**).

DNA extraction methods revealed kit biases that affected relative abundance measurements for the starting consortium material. Discrepancies between DNA extraction kits were notable as they painted different pictures about the key guilds required for NH_3_ removal. Members of the *Nitrosospira* are known AOB that rely on handoffs to NOB including members of the *Nitrobacter* genus to complete the oxidation of NH_3_ to nitrate (NO_3_^-^) (26, 27). Previous work on the physiology of these taxa aligns with our functional characterization of the consortium in development. MAGs from these genera carried key genes for NH_3_ and NO_2_^-^ oxidation (see **Figure 1**). Characterizing the consortium based on the Metagenom Bio kit showed dominance by an AOB *Nitrosospira* genus, whereas the Qiagen Powersoil kit showed members of the *Nitrosospira* occurring at comparable relative abundance to NOB from the *Nitrobacter* genus (see **Figure S1** and **Supporting Results**). Although it is unclear which kit provides a true representation of the composition, this finding highlights the importance of a holistic approach to characterize microbial consortia.

**Figure 1:**
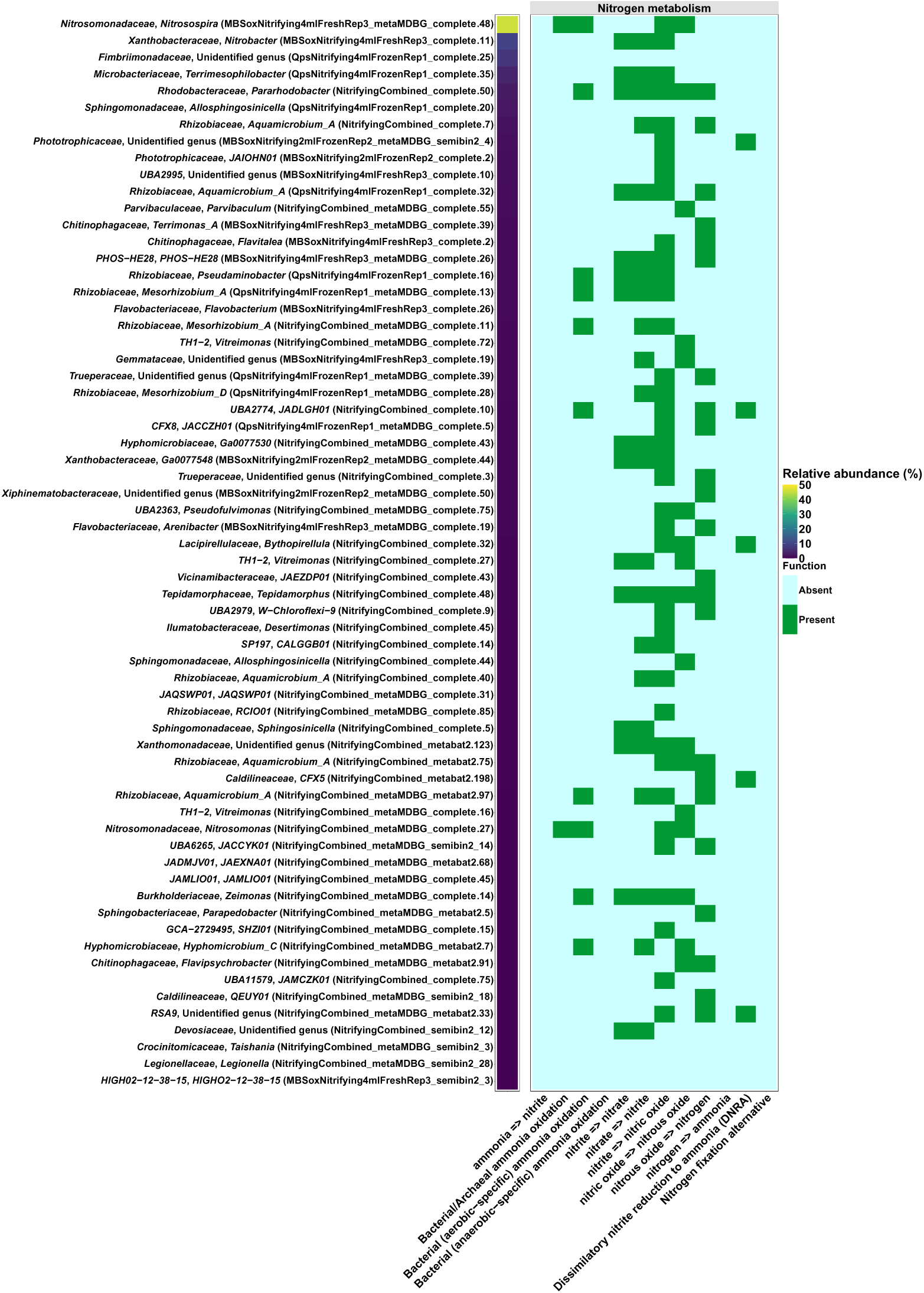
Relative abundance and nitrogen metabolism for 64 metagenome assembled genomes (MAGs) built using PacBio long-read sequencing. Relative abundance data was generated from the combined reads of multiple DNA extraction methods that were used to assemble MAGs.

Considering the differences observed for relative abundance between extraction methods, we combined the entire read sets in subsequent relative abundance analyses of the consortium starting material. MAG MBSoxNitrifying4mlFreshRep3_metaMDBG_complete.48 classified to the genus *Nitrosospira* (herein referred to as the dominant *Nitrosospira* population) maintained a high relative abundance of 45.96% whereas MBSoxNitrifying4mLFreshRep3_complete.11 classified to the *Nitrobacter* genus had a relative abundance of 9.73% in the combined dataset (see **Figure 1**). Beyond the dominant population of *Nitrosospira*, 11 MAGs displayed relative abundance values between 1 and 10 % and the remaining 52 MAGs were < 1 % abundance (see **Figure 1** and sheet #1 in **Supplementary Data 1**). The abundance of the dominant *Nitrosospira* aligns with the expected composition of this culture, which is maintained under conditions that favour AOB (see **Methods**). The lower abundance of the NOB *Nitrobacter* population suggests that key guilds required for nitrification do not occur at a 1:1 ratio, although this is difficult to disentangle from the kit biases noted above. Outside of the dominant *Nitrosospira* population, only MAG NitrifyingCombined_metaMDBG_complete.27 classified to the genus *Nitrosomonas* displayed the metabolic capacity for NH_3_ oxidation (see **Figure 1**). This MAG occurred at low relative abundance of 0.15% suggesting it is not a major contributor to nitrification (see sheet #1 in **Supplementary Data 1**). No genera with the potential for the complete oxidation of NH_3_ (*i.e.,* commamox) or the anaerobic NH_3_ oxidation (*i.e.,* annamox) were detected in the starting material (see **Figure 1).**

While NH_3_ oxidation was most apparent in the dominant population of *Nitrosospira,* 9 MAGs from the families *Burkholderiaceae*, *Hyphomicrobiaceae*, *Rhizobiaceae*, *Rhobacteraceae*, and the unnamed family UBA2774 carried genes for hydroxylamine oxidation to NO_2_^-^ based on functional analyses (see **Figure 1**). These populations occurred at an order of magnitude lower relative abundance (ca. 0.12 % to 3.08%) compared to the dominant *Nitrosospira* population suggesting they are less important contributors to nitrogen cycling overall in the consortium (see **Figure 1**). Interestingly, many of these MAGs also carried the genes for NO_2_^-^ oxidation suggesting they could produce NO_3_^-^ following NH_3_ oxidation by members of the *Nitrosospira* population (see **Figure 1**). There were 14 other MAGs classified to the families *Burkholderiaceae*, *Devosiaceae*, *Hyphomicrobiaceae*, *Microbacteriaceae*, PHOS-HE28, *Rhizobiaceae*, *Sphingomonadaceae*, *Tepidamorphaceae*, TH1-2, *Xanthobacteraceae*, and *Xanthomonadaceae* that displayed the potential to oxidize NO_2_^-^ to NO_3_^-^ (see **Figure 1**). These MAGs occurred at low relative abundance values < 5 % and carried genes for denitrification suggesting oxidized nitrogen species could also be removed through the production of N_2_ (see **Figure 1**).

None of the MAGs in the starting material carried genes for nitrogen fixation, which could potentially resupply NH_3_ in a setting in contact with the atmosphere (see **Figure 1**). Five MAGs with relative abundance < 1.5 % classified to unnamed genera in the families *Caldinilneaceae*, *Lacipirellulaceae*, *Phototriphicaceae*, RSA9, and UBA2774 carried the genes for dissimilatory NO_2_^-^ reduction, which could counteract nitrification activity (see **Figure 1**). Further analysis using DRAM identified five MAGs carrying the gene coding for the assimilatory NO_3_^-^ reductase catalytic subunit (*i.e.,* KEGG ID K00372) (see sheet #2 in **Supplementary Data 1**). These MAGs spanned the families JACCZH01, *Hyphomicrobiaceae, Rhodobacteraceae*, and *Xanthobacteraceae* at abundances < 3%, suggesting assimilatory NO_3_^-^ reducers are less favoured in the consortium.

Overall, the taxonomic and putative nitrogen metabolism characterization of the consortium using long-read sequencing aligns with its intended application in WWTPs. Genera containing AOB and NOB occurred at high abundances, suggesting the target guilds are positively selected for during cultivation. Deep long-read DNA sequencing also led to the recovery of lower abundance MAGs that potentially compete for nitrogen substrates alongside AOB and NOB via denitrification, dissimilatory NO_3_^-^ reduction, and assimilatory NO_3_^-^ reduction. Despite many of these MAGs having between 99 and 100% completion, we were unable to classify most MAGs to the species level. We attribute the lack of species resolution to members of this consortium being poorly represented in GTDB despite a number of studies characterizing communities in WWTP settings using metagenomic techniques (23, 24, 28–30). Improving the representation of microbial taxa from engineered systems where commercial consortia can be sourced through top-down approaches will be important to leveraging multi-omics strategies that provide compositional data that can be used as part of risk assessment.

Increasing this representation will also be important for assessing potential human health threats tied to commercial microbial products. Although we didn’t explicitly search for putative pathogens, long-read sequencing led the detection of a MAG classified to an unnamed species in the *Legionella* genus (*i.e.*, NitrifyingCombined_metaMDBG_semibin2_28 – 87% complete), which includes known pathogens associated with WWTPs (31, 32). A full analyses of virulence genes and other markers of pathogenicity is outside the scope of this work. We make note of this observation to further demonstrate the value of multi-omics strategies for assessing human health risks tied to commercially available microbial consortia.

### Cell growth and nitrogen cycling along a redox gradient

We conducted growth experiments for a total of eight weeks in line with the medium refreshment schedule for this consortium (see **Methods**) to compare how the microbial community responsible for nitrogen cycling was affected by a redox gradient representative of WWTPs. All bottles that received live consortium material showed a 3 to 4-fold increase in O.D. 600 nm over the course of the experiment up to ∼0.4 (see **Figure S2**). No growth was observed in the sterile controls except for replicate #2 in the nitrate reduction treatment, which was likely contaminated around day 30 (see **Figure S2**). Aerobic and anaerobic cultures initially experienced a decrease in pH to 6.0 and only aerobic cultures saw a return to circumneutral pH values of 7.5 during the experiment (see **Figure S2**). These observations suggest that despite the manufacturer maintaining oxic conditions to select for AOB, the consortium in development can grow under anoxic conditions.

Abiotic nitrogen cycling was not observed in sterile controls. Colorimetric NH_3_ measurements on day 0 were higher than the theoretical total nitrogen in the synthetic pondwater (*i.e.,* 5.0 to 7.5 mM vs 3.11 to 4.11 mM) but stabilized to values in line with expectations afterwards (*i.e.,* 0 to 2.7 mM) (see **Figure 2**). NO_2_^-^ concentrations remained stable in abiotic controls and in line with the expected composition of the medium (ca. 0.47 mM to 0.82 mM) except for replicate #2 from the nitrate reduction treatment, which was contaminated (see **Figure 2**). NO_3_^-^ concentrations were stable in abiotic controls ranging from 0.19 to 0.21 mM, which were slightly lower than the theoretical 0.84 mM added to the medium (see **Figure 2**). We discuss sources of analytical variance for the nitrogen chemistry data in more detail in the *Nitrogen Chemical Analyses* section of the **Supporting Results**. Given that nitrogen measurements for all bottles were carried out identically and subject to the same limitations, we deem it suitable to compare nitrogen cycling over time through a phenotypic lens to support our microbiological analyses.

**Figure 2:**
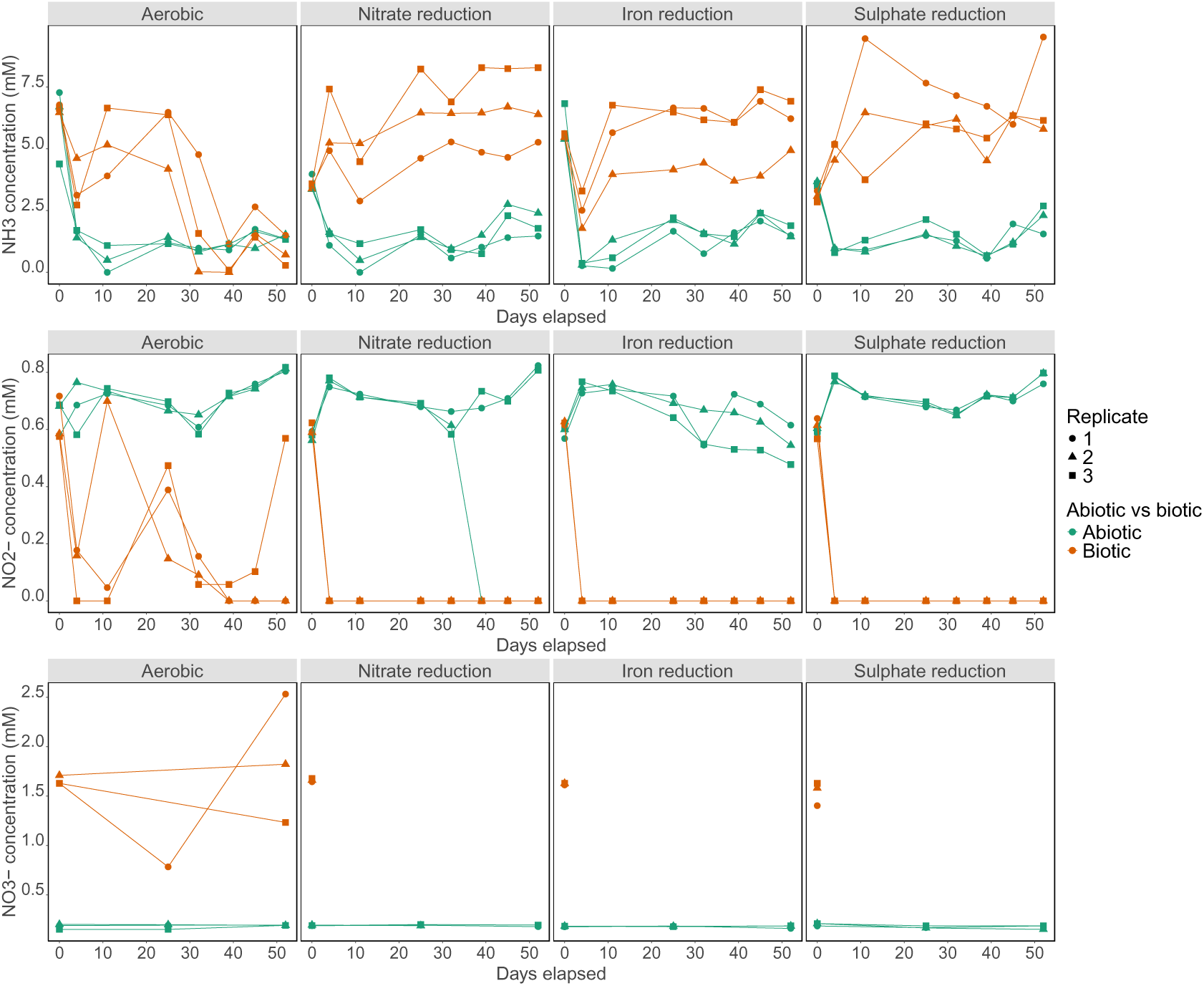
NH_3_, NO_2_^-^ and NO_3_^-^ concentrations for enrichment cultures inoculated with starting material from the nitrifying consortium in development and sterile controls grown along a redox gradient in synthetic pond water. NH_3_ and NO_2_^-^ were measured frequently during the incubation period using colorimetric methods whereas NO_3_^-^ analyses could only be carried out for select samples tied to sequencing efforts. The aerobic cultures were purged daily with oxygen and anaerobic cultures were maintained under a headspace of 97% N_2_/3% H_2_ and handled in an anaerobic glovebox. The nitrate reduction treatment was grown with the NO_3_^-^ already present in the medium acting as an electron acceptor whereas the iron reduction and sulphate reduction treatments were amended with Fe^III^ and SO_4_^2-^ as terminal electron acceptors, respectively. Abiotic vs biotic treatments are colour coded and triplicates are denoted by different shapes.

Live cultures differed considerably in the nitrogen cycling observed compared to abiotic controls. Aerobic cultures did not completely oxidize NH_3_ in the beginning of the experiment and concentrations fluctuated considerably over the ensuing weeks. Aerobic cultures initially decreased NH_3_ concentrations from ∼ 7.0 mM to 2.5 to 4.5 mM in the first seven days, however NH_3_ concentrations increased up to 5.0 to 7.5 mM between days 10 and 20 with considerable variance between replicates (see **Figure 2**). Additional decreases in NH_3_ concentrations below 2.5 mM occurred after 30 days in all aerobic cultures suggesting a considerable lag period prior to net NH_3_ removal occurring (see **Figure 2**). The initial increases in NH_3_ in aerobic cultures were accompanied by fluctuations in NO_2_^-^ concentrations from 0.2 to 0.8 mM, however the detection of 1.5 to 2.5 mM NO_3_^-^, the product of nitrification, did not occur until after day 30 when NH_3_ removal occurred more consistently (see **Figure 2**). In contrast to aerobic cultures, anaerobic cultures saw the total loss of NO_2_^-^ and NO_3_^-^ alongside plateaus in NH_3_ concentrations that remained stable between 5.0 to 7.5 mM over the time frame of the experiment (see **Figure 2**). This trend was consistent regardless of the terminal electron acceptor supplied.

The pronounced lag in NH_3_ removal observed for aerobic cultures was unexpected. We attribute this lag to suboptimal growth conditions encountered in the aerobic metabolic treatment. Our choice to supply oxygen periodically rather than continuously (see **Methods**) potentially limited the oxygen required for NH_3_ oxidation and aerobic respiration. Although it is possible that some nitrification occurred over the first week of incubating the aerobic cultures, it is also possible that decreases in pH led to inhibition (see **Figure S2**). The manufacturer of the consortium in development indicated that aerobic nitrification is inhibited at pH < 6.8 (personal communication), which is supported by other physiological studies on AOB where the lower pH favours the NH_4_^+^ ion vs the NH_3_ compatible with the AmoA protein responsible for oxidation (33). All aerobic cultures experienced pH decreases < 6.8 within the first week of the experiment prior to pH returning to ∼7-8, which may have prolonged the lag phase for nitrifiers (see **Figure S2**). Regardless of the mechanisms that contributed to the lag period, these observations highlight the potential for the consortium to shift towards NH_3_ production, rather than removal, under oxygen limitations and time frames representative of WWTP settings. We examine these observations through a functional lens using metagenomic and metatranscriptomic data in the following section.

### Shifts in taxonomic composition and nitrogen cycling in response to a redox gradient

Short-read metagenomic and metatranscriptomic sequencing were used to demonstrate how increasingly affordable meta-omics techniques could be used by stakeholders to assess shifts in composition and function for commercial microbial consortia. Although we successfully applied long-read sequencing to the starting consortium material, short-read sequencing was selected to circumvent the need for higher DNA yields in long read sequencing. We were unable to obtain sufficient yields due to the low cellular growth observed during the incubation (see **Figure S2**).

Beta diversity analysis using non-metric multidimensional scaling analysis (NMDS) showed that anaerobic cultures maintained a similar composition as the starting inoculum (see **Figure 3**). Aerobic cultures diverged in their composition relative to the inoculum and the anaerobic cultures (see **Figure 3**). Data points for aerobic cultures from week 4 clustered away from the inoculum but also each other (see **Figure 3**). The composition of aerobic replicates #1 and #2 overlapped at week 8, suggesting these cultures converged towards a similar composition whereas replicate #3 clustered closer to samples obtained at week 4 (see **Figure 3**). These observations suggest that aerobic replicates did not experience consistent shifts in community composition despite being established with the same starting biomass. When we omitted aerobic cultures from the NMDS analysis, differences between anaerobic cultures and the inoculum were more apparent but the degree of divergence varied depending on the time point (see **Figure S3**). These observations show that minor differences in composition occurred between the starting material and anaerobic cultures compared to the aerobic cultures.

**Figure 3:**
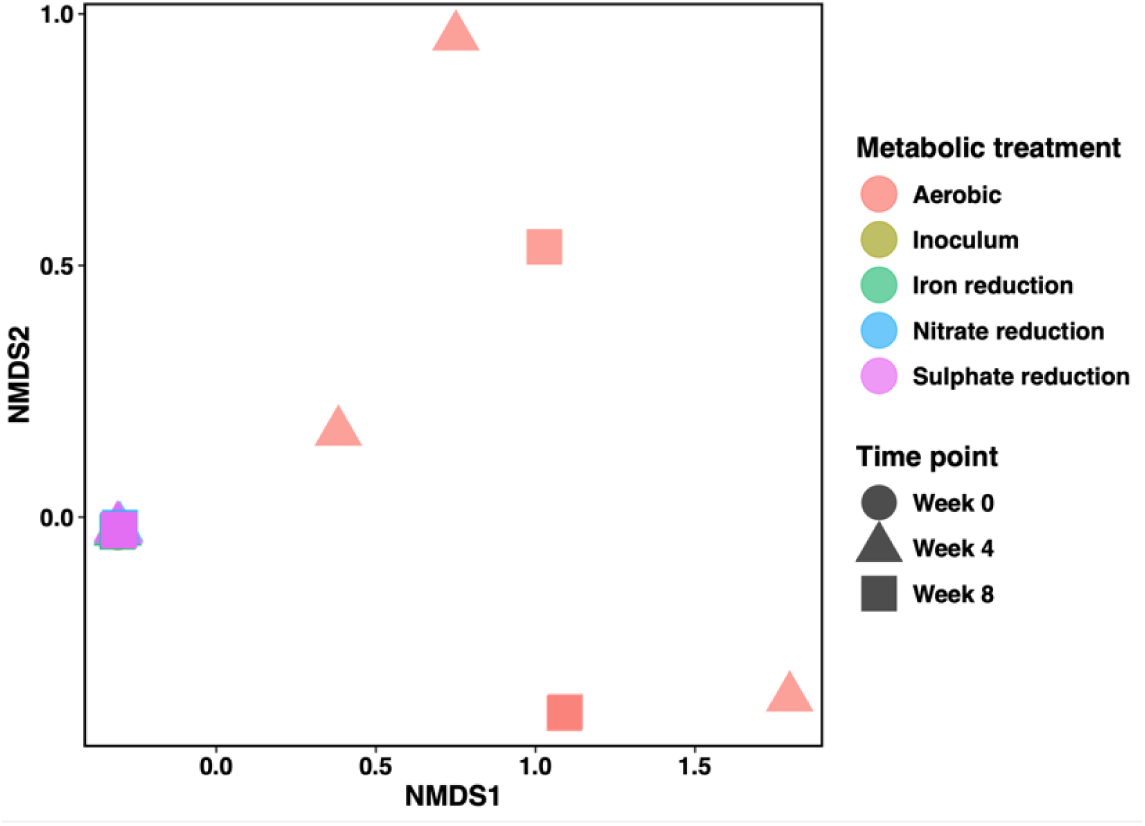
Non-metric multidimensional scaling (NMDS) based on the relative abundance of 98 metagenome-assembled-genomes (MAGs) recovered from enrichment cultures grown along a redox gradient for eight weeks. The Bray-Curtis dissimilarity index was used to generate a distance matrix for NMDS. The default commands ‘metaMDS’ from the ‘vegan’ v. 2.6.4 package was used to run 20 iterations of the NMDS ordination, which provided a stress value of 0.0279. Metabolic treatments and the inoculum material have been colour-coded and times points are identified by different shapes.

The divergence in overall community composition was likely driven by shifts in the abundance of key guilds involved in nitrification. Aerobic cultures saw a decrease in the relative abundance of the AOB *Nitrosospira* population that initially dominated the inoculum (*i.e.,* MAG MBSoxNitrifying4mlFreshRep3_metaMDBG_complete.48). This MAG was detected at relative abundances of ∼ 65 to 70% at Week 0 in the starting material and decreased to a range of 3.10 to 26.42 % across aerobic replicates as the incubation progressed (see **Figure 4**). Notably, replicate #2 from the aerobic cultures maintained the highest abundance of *Nitrosospira* at 26.42% for week 4 (see **Figure 4**). This sample was obtained just after a peak in NO_2_^-^ was measured alongside NH_3_ loss and the *amoABC* genes required for nitrification were upregulated according to RNA read counts, suggesting the *Nitrosospira* population was likely contributing to nitrification after the lag (see **Figure 2, Figure S4**, and **Supporting Results**). Relative abundance values for the *Nitrosospira* population stabilized between 11.60 to 17.43 % at week 8 when NH_3_ concentrations decreased in all three aerobic cultures (see **Figure 2** and **Figure 4**). The stable decrease in *Nitrosospira* abundance was accompanied by decreases in abundance of the NOB genus *Nitrobacter* (*i.e.,* MAG MBSoxNitrifying4mlFreshRep3_complete.11) from ∼10% to < 2% and lower transcription of genes required for nitrification at week 8 (see **Figure 4**, **Figure S4** and **Supporting Results**). These observations suggest that substrate limitations inhibited the growth of key nitrification guilds towards the end of the cultivation experiment.

**Figure 4:**
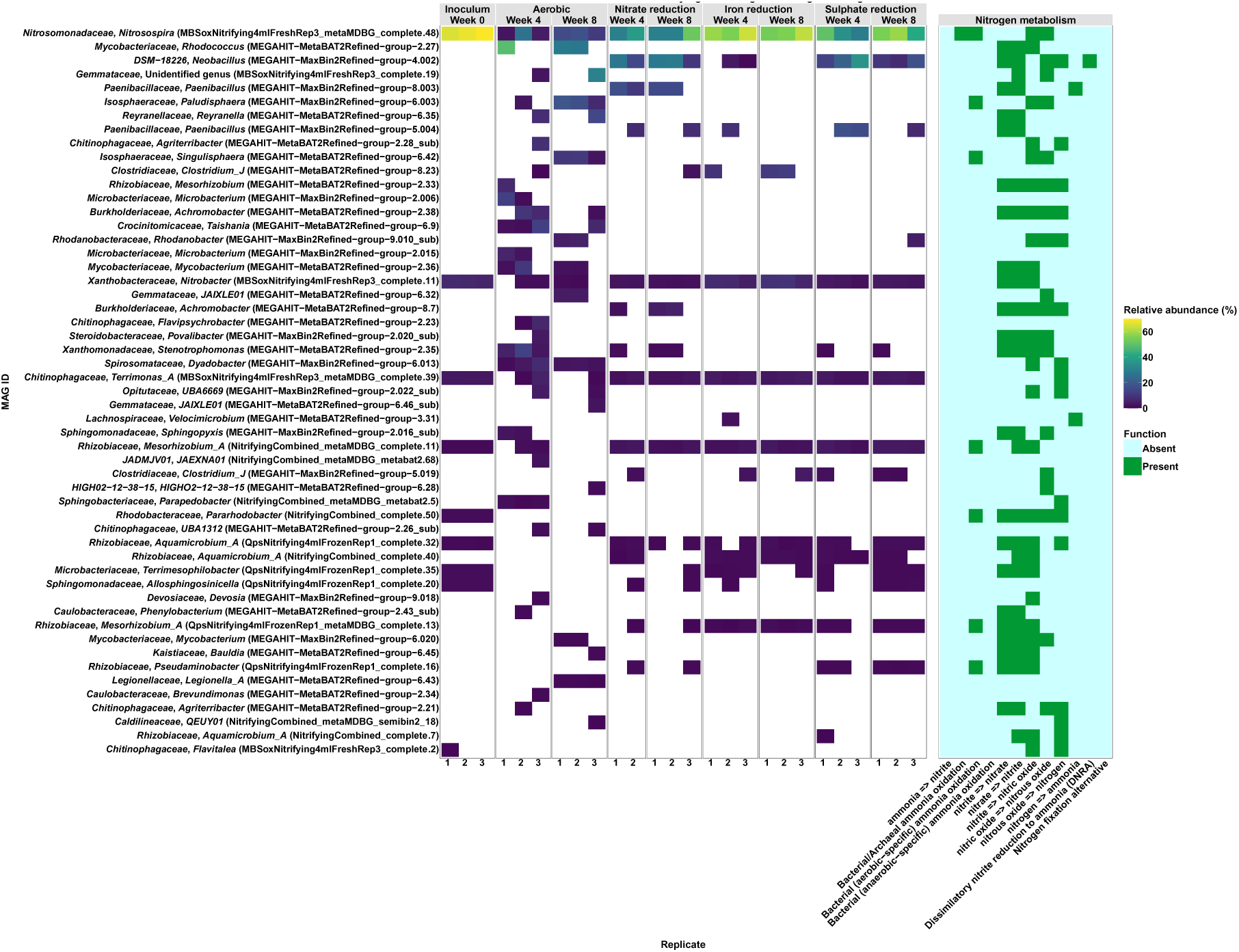
Relative abundance for metagenome-assembled genomes (MAGs) >= to 1 % and nitrogen metabolism for enrichment cultures established with the nitrifying consortium in development as starting material and grown along a redox gradient for eight weeks. Enrichment culture samples were exclusively sequenced using Illumina shotgun metagenomic sequencing. All MAGs assembled from the starting material and enrichment culture time series were combined into a final dereplicated series of MAGs to assess relative abundance.

Major shifts in the dominant AOB population of *Nitrosospira* were not apparent in the anaerobic cultures. Most anaerobic cultures maintained *Nitrosospira* populations at relative abundances that varied between 29.10 to 64.53%, which is comparable to the 65 to 70% observed at week 0 (see **Figure 4**). The NOB *Nitrobacter* population also occurred at relative abundances between 2.18 to 9.61%, which were comparable to the starting material (see **Figure 4**). These observations suggest that the onset of anoxic conditions did not significantly affect growth of the key populations required for nitrification, which was unexpected given the constant aeration conditions used by the manufacturer to favour these guilds.

In this instance, we attribute the maintenance of high abundances of the *Nitrosospira* populations in anaerobic cultures to a shift towards anaerobic metabolism. Functional analyses of *Nitrosospira* MAGs showed these populations have the capacity for NO_2_^-^ reduction to nitric oxide (NO) and NO reduction to nitrous oxide (N_2_O) (see **Figure 1** and **Figure 4**). These populations occurred alongside a variety of other taxa in the starting material that had the capacity to reduce NO to nitrogen gas (N_2_) to complete the denitrification process (see **Figure 4**). RNA read counts showed an increase in the abundance of the denitrification KEGG orthologs K00368 coding the NO-forming NO_2_^-^ reductase and K00376 coding the N_2_O reductase in anaerobic cultures when compared to aerobic cultures at the same time points (see **Figure S4**). These findings show there are increases in the transcription of anaerobic nitrogen redox cycling machinery that could fuel the anaerobic growth observed in these cultures (see **Figure S2**).

Several MAGs with the capacity for nitrogen redox cycling that were not detected in the inoculum material at week 0 were detected in weeks 4 and 8 whereas many rare taxa that occurred below 1% abundance at week 0 were absent in aerobic cultures during the same time (see **Figure S5**). We suspect that the shifts in the most abundant nitrogen cycling taxa alongside the disappearance of rare taxa are driving differences in beta diversity noted previously (see **Figure 3**). Many of these newly detected MAGs occurred at relative abundance values between 10 to 35 %, suggesting they were important members of the community that were enriched via the different metabolic treatments applied (see **Figure 4**).

An example of one such population in the aerobic cultures is represented by MAG MEGAHIT−MetaBAT2Refined−group−2.27 classified to *Rhodococcus* sp00434560, which carried genes for NO_2_ ^-^ oxidation and partial denitrification. This *Rhodocococcus* population maintained a relative abundance ranging from 27.78% (week 8, replicate 2) to 48.82% (week 4, replicate 1) throughout the experiment (see **Figure 4**). Similarly, MAG MEGAHIT−MaxBin2Refined−group−6.003 classified to an unidentified species in the *Paludisphaera* genus carried genes for hydroxylamine oxidation and partial denitrification and maintained relative abundances between 7.59 and 18.82 % by week 8 in aerobic cultures (see **Figure 4**). We noted an increase in the proportions of RNA reads mapped to MAG MEGAHIT−MetaBAT2Refined−group−2.27 from the *Rhodococcus* genus (ca. 34%) and a lesser increase of the RNA mapped to MEGAHIT−MaxBin2Refined−group−6.003 from the *Paludisphaera* genus (ca. 2 to 5%) corresponding to these time points (see **Figure S6**). This observation suggests these populations also contributed significantly to metabolic activity in these cultures although this is difficult to fully contextualize given that a large proportion of RNA reads did not map to MAGs assembled from these cultures (see **Figure S6**).

MAGs with nitrogen cycling capabilities that precluded detection in the starting material at week 0 were detected at weeks 4 and 8 in anaerobic cultures. We suspect the combination of enrichment of select MAGs alongside the varied increases of low abundance taxa are driving compositional differences between anaerobic cultures and the starting material (see **Figure S3**). Anaerobic cultures had increases in abundance for MAG MEGAHIT−MaxBin2Refined−group−4.002 classified to an unidentified *Neobacillus* species with the capacity for near complete denitrification, although this varied considerably across treatments (ca. from 0.19 % to 35 %) (see **Figure 4**). Notably, this *Neobacillus* population was one of the few MAGs carrying genes for dissimilatory NO_3_^-^ reduction to NH_3_ (DNRA), which could be contributing to the NH_3_ plateaus observed in anaerobic treatments (see **Figure 2** and **Figure 4**). MAG MEGAHIT−MaxBin2Refined−group−8.003 classified to an unidentified species in the genus *Paenibacillus* also had the capacity oxidize NO_2_^-^ to NO_3_^-^ and *vice versa* and occurred at relative abundances between 11.10 to 17.01 % in the nitrate reduction treatment (see **Figure 4**). This *Paenibacillus* population had nitrogen fixation capacity, highlighting another potential pathway that could resupply NH_3_ under anoxia (see **Figure 4**).

These populations coexisted with other MAGs with low abundance (∼ 1% or lower) from several different genera (see **Figure S5**). The emergence of these new populations aligned with variable increases in the proportion of RNA mapped to MAG MEGAHIT−MaxBin2Refined−group−4.002 from the *Neobacillus* genus (reaching > 70 % in some cases) and lesser increases in RNA mapped to the MEGAHIT−MaxBin2Refined−group−8.003 (ca. 3 to 5%) from the *Paenibacillus* genus (see **Figure S6**). The MAG MEGAHIT-MaxBin2Refined−group 5.004, which was also classified to the genus *Paenibacillus* and had similar metabolic capacity to MEGAHIT−MaxBin2Refined−group−8.003, accounted for > 20 % of the RNA reads mapped to MAG in some cases despite being sporadically detected at abundances ranging from <1 % up to 17.42% in some cultures (see **Figure 4** and **Figure S6**). The increased RNA mapped to these bins occurred alongside increases in transcription for denitrification genes suggesting these populations were contributing to nitrogen cycling via those pathways in anaerobic cultures (see **Figure S4** and **Supporting Results**). These observations further suggest different *Paenibacillus* populations that could not be detected in the inoculum were selected for under anoxic conditions.

Although specific populations were detected in anaerobic cultures that could contribute to NH_3_ through assimilatory and nitrogen fixation pathways, there was no evidence of these pathways in aerobic cultures that saw comparable increases in NH_3_ concentrations for the first four weeks of the incubation (see **Figure 4**). Instead, we suspect that aerobic cultures were able to access undefined sources of NH_3_ contained in the yeast extract that was supplied to mimic complex nutrients typically found in WWTPs (34). When triaging metatranscriptomics results to the top 25 values obtained for RNA read counts associated with KEGG IDs for amino acid metabolic pathways, we observed increased transcriptions for genes coding for glutamate dehydrogenase and glutamine synthetase that can catalyze deaminating reactions to produce NH_3_ in aerobic cultures (see **Figure S7** and **Supporting Results**). These analyses also showed that anaerobic cultures had increased transcription of genes coding enzymes capable of deamination including glutamine synthetase and glutamate synthase that could also contribute to NH_3_ alongside some of the assimilatory and nitrogen fixation pathways noted previously (see **Figure S7**). These findings underscore the value of using a multi-omic approach to explore other metabolic pathways that could affect the performance of a commercial microbial consortia when faced with realistic but suboptimal conditions in less controlled settings.

### Conclusions

In this study, we successfully applied long-read DNA and short-read DNA and RNA sequencing to characterize a consortium in development for wastewater treatment. We demonstrated that long-read sequencing enabled the recovery of nearly complete genomes from microbes whose functions aligned with the intended use of this consortium for NH_3_ removal. Stakeholders should consider a comprehensive approach incorporating multiple DNA extraction methods to account for bias when characterizing the composition of microbial mixtures and consortia for quality assurance and regulatory purposes. We showed meta-omic approaches can be used to monitor shifts in the composition and desired performance of this model consortium under conditions representative of a WWTP. Whole-community techniques revealed that taxa not initially detected in starting material could become abundant and metabolically active under different growth conditions with significant impacts on nitrogen cycling. These multi-omic analyses informed a conceptual explanation wherein metabolically versatile nitrogen redox cycling bacteria shifted away from aerobic nitrification in favour of denitrification and deaminating pathways leading to lags in NH_3_ removal when oxygen was provided and net NH_3_ production under oxygen limitation.

This work demonstrates how regulators could leverage multi-omics approaches in the risk assessment of commercially available microbial consortia. Despite using long-read and short-read metagenomic sequencing for taxonomic classification, we were unable to identify many taxa in this consortium to a species level. This finding highlights why carrying out risk assessments based on compositional data may be challenging and how functional analyses can fill critical gaps in our understanding of desirable and undesirable consortia functions. Although it is unrealistic for stakeholders to try and capture how these functions may shift along an indeterminate combination of environmental variables, our work highlights how even small-scale controlled cultivation experiments can provide valuable mechanistic information for predicting consortia performance *in situ*.

In the future, it will be crucial to refine and validate a framework with other available consortia products to ensure reproducibility across microbial products with different metabolic functions. Although multi-omics methods must be tailored to specific analytical endpoints of interest, ranging from contaminant removal, natural product synthesis, or assessing the capacity for virulence, this study provides a proof of principle demonstrating how empirical multi-omics data can support evidence-based decision making. Supporting regulators in shifting the paradigm on how these products are managed will require developing a logical framework on how to standardize and disseminate the analyses of such complex multi-omics data to support the decision-making process. Developing these frameworks with input from manufacturers and end users in different biotechnology sectors will be important to ensuring uptake and compliance. This will be a critical step to effectively translating multi-omics studies into tangible policy actions that will ensure the safe and effective of use of microbial products that have undergone assessment.

## Methods

### Initial consortium maintenance

This work was carried out in collaboration with an industry partner that has requested anonymity. The partner kindly provided a consortium that was in development for removing NH_3_ from wastewater. This culture was maintained in a proprietary defined medium under constant aeration. The medium was refreshed every six to eight weeks, which informed the timeframes applied in our growth experiments. All other information related to culture maintenance is proprietary.

### Long-read PacBio sequencing on the starting consortium material

A single bottle containing ∼ 1 L of culture of the nitrifying consortium in development was shipped at room temperature and stored at 4°C after receiving. Aliquots were frozen in sterile 15 mL conical tubes and stored at -80°C to preserve the culture profile with 4 mL of thawed sample being used for extraction or 2 mL of fresh culture. Cell pellets were extracted using the PowerSoil Pro kit (Qiagen) and the Sox DNA Extraction kit (Metagenom Bio Life Science). 4 µL of RNase A (100mg/ml) was added after each kit’s bead beating step and tubes were incubated for five minutes at room temperature to reduce RNA that may interfere with PacBio sequencing. Samples were purified and concentrated with the Genomic DNA Clean & Concentrator^TM^-10 Kit (Zymo Research). PacBio Sequel II sequencing was performed at Integrated Microbiome Research (Dalhousie University, Halifax, NS) using the SMRTbell Prep Kit 3.0 for shotgun metagenomics. The samples were already sheared sufficiently for library preparation from the extractions and 1.82 million (22.6 Gbp) total reads were generated from three samples from the two extraction kits.

### Enrichment experiment setup

A fresh sample from the same lot of the nitrifying consortium in development previously sequenced by PacBio was used to inoculate enrichment cultures that were grown in a synthetic pond water recipe provided by the consortium manufacturer. Synthetic pondwater was prepared by dissolving the ingredients listed in **Table 1** in 1 L of ultra-filtered water (*i.e.,* Milli-Q) prior to adjusting pH to 7 with 0.5 mL of 1 M NaOH and autoclaving at 121°C for 30 min.

**Table 1:** Synthetic pondwater recipe.

| Medium component | Reagent | Concentration (g L<sup>-1</sup>) | Concentration (mM) |
| --- | --- | --- | --- |
| Basal medium | KNO<sub>3</sub> | 0.085 | 0.84 |
|  | NaNO<sub>2</sub> | 0.06 | 0.87 |
|  | NH<sub>4</sub>Cl | 0.075 | 1.4 |
|  | H<sub>3</sub>PO<sub>4</sub> | 0.038 | 0.39 |
|  | Yeast extract | 1 | NA |
|  | Glucose | 1 | 5.55 |
| Fe<sup>III</sup> stock (1 L) 100X<br>(10 mL in 1 L medium) | Fe<sup>III</sup>-ammonium-citrate<sup>d</sup> | 0.262 | 1 |
| SO<sub>4</sub><sup>2-</sup> stock (1 L) 100X<br>(10 mL in 1 L medium) | Na<sub>2</sub>SO<sub>4</sub> | 0.142 | 1 |

We prepared cultures and abiotic controls for the different metabolic in triplicate by filling 90 or 100 mL of medium, respectively, into 150 ml serum bottles sealed with butyl rubber stoppers that were crimped shut. Aerobic cultures were filled with basal medium stored on the bench and purged daily on working days, Monday to Friday, with a sterile supply of atmospheric air that passed through an autoclaved 0.22 µm pore-size PTFE filter. We chose to grow aerobic cultures in this way rather than in containers open to the atmosphere that were shaken to ensure consistent containers and sampling equipment was used for all cultures in this experiment. The medium used for anaerobic treatments was bubbled under a sterile flow of N_2_ for 30 minutes prior to being moved to a Coy anaerobic glovebox that maintained an atmosphere of 97% N_2_/3% H_2_ for aliquoting into serum bottles conditioned under anoxia for 48 hours. The anaerobic metabolic treatment for nitrate (NO_3_^-^) reduction was not amended with any other reagents because NO_3_ ^-^ was already present in the medium (see **Table 1**). Metabolic treatments for iron reduction and sulphate reduction were amended to a final concentration of 1 mM of Fe^III^ or SO_4_^2-^ from 100-fold stocks that were filter-sterilized and bubbled under N_2_ prior to use (see **Table 1**).

Sterile abiotic controls did not receive any material for inoculum. Biotic samples were inoculated at 10% (v/v) using biomass from a freshly received batch of the consortium using sterile 10 mL syringes. Inoculum material for aerobic cultures were handled on the bench and the same material was subsequently bubbled under sterile N_2_ prior to inoculating anaerobic cultures to minimize oxygen intrusion. A 5 mL subsample was withdrawn from the starting material in addition to abiotic and biotic bottles prior to being filtered through a 0.22 µm pore size PES filter, measured for pH and redox potential, and stored at -20°C until nitrogen chemistry analysis. 2 mL aliquots were also obtained in triplicate for RNA extraction. These samples were placed in sterile conical tubes and topped up with 10 mL of DNA/RNA shield (Zymo Research) and stored at -20°C until processing. Additional 2 mL aliquots were withdrawn in triplicate from the starting material for DNA acid extractions. These samples were centrifuged at a speed of 10,000 g for 8 minutes prior to decanting the supernatant and storing pellets at -80°C until processing.

Abiotic and biotic bottles had 5 mL withdrawn weekly for nitrogen chemistry, pH, and ORP measurements as outlined above. Additional 0.2 mL aliquots were also retrieved to measure cell growth at O.D. 600 nm in a 96-well plate using a plate reader with a monochromator. Subsamples for nucleic acid extraction were withdrawn bi-weekly and stored as noted above.

### Nitrogen chemistry analyses

NO_2_^-^ and NH_3_ were measured using dedicated colorimetric methods adapted from the literature. A select number of subsamples were sent to an independent certified lab (i.e., Bureau Veritas, ISO 17025) to carry out NO_2_^-^ and NO_3_^-^ analyses for select archived time points that were representative of the beginning, middle, and end of the experiment using the SM 23 4500-NO_3_I/NO_2_B protocol. NO_2_^-^ analyses were conducted using a modified Griess-reagent based method (Invitrogen, G7921). The Griess reagent was prepared by mixing equal parts of N-(1-naphthyl)ethylenediamine and sulfanilic acid. Samples were thawed daily and diluted 10-fold in a modified synthetic pondwater where all defined nitrogen compounds were removed to control for potential background coming from undefined sources of nitrogen such as yeast extract. Standards were created daily to establish calibration curves ranging from 0.08 mM up to 1 mM, which we established as the linear range of the assay with the equipment available. To prepare a sample for NO_2_^-^ analyses, 86.6 µL of Milli-Q water and 40 µL of the Griess reagent was added to target wells in a 96-well plate. 100 µL of the desired subsample was then added to the well and mixed by pipetting up and down prior to incubating at room temperature for 30 minutes and reading the absorbance on a plate reader at O.D. 584 nm. All samples were blank corrected according to their corresponding medium recipe that included the different terminal electron acceptors added to the medium (see **Table 1**).

Total NH_3_ was measured using a miniaturized version of the indophenol colorimetric method adapted for a 96-well microplate reader (35). Note that in the original method paper, the authors refer to NH_3_ and ammonium (NH_4_^+^) interchangeably to refer to the nitrogen species. In this same paper, the authors do indicate that NH_4_^+^ would dominate speciation in systems with pH < 9 such as ours due to its pKa value of 9.25 (35). In this study, we chose to report the results as NH_3_ because the original citation did not distinguish between the two nitrogen species and to ensure our reporting is consistent with literature on AOB, which require NH_3_ as a substrate (36, 37). Standards were prepared in the same background synthetic pond water to control for the pH sensitivity of the colorimetric assay and any undefined sources of nitrogen originating from the yeast extract in the growth medium. Experimental samples needed to undergo a 10-fold dilution in synthetic pond water devoid of defined nitrogen sources to ensure they fell within the linear range of the assay in most cases (see **Table S1** and **Supporting Results**). A master mix was created based on the number of tubes (n) that needed to be analyzed where n x 30 µL sodium nitroprusside and n x 75 µL of oxidizing solution (hypochlorite) were used. 105 µL of this master mix was aliquoted into 2 mL tubes followed by the addition of 750 μL of sample or standard. Following this step, 30 µL of phenol solution was added to each tube and gently mixed by pipetting before transferring each sample to 96-well plate. Plates were incubated in the dark at room temperature for 60 min to allow the colour to develop before measuring O.D. at 640 nm on a plate reader.

### Short-read metagenomic and metatranscriptomic sequencing on enrichment cultures

Suboptimal RNA yields for library preparation were recovered from samples preserved in DNA/RNA shield and so pellets preserved for DNA extraction were subsequently tested and used for DNA/RNA co-elution for short-read sequencing. Samples were extracted from 2 mL of pelleted cells using the Zymo DNA/RNA Microbiome kit to co-elute each fraction from the same sample. Bead beating homogenization was completed on a VWR Mini beater at 5S speed for 70 seconds repeated three times in five-minute intervals. DNA samples were purified and concentrated as previously described but were treated with 4 uL of RNAse A (100 mg/ml) for 10 minutes at room temperature before purification. RNA was treated with DNase I (Zymo Research) and purified using the RNA Clean and Concentrator Kit^TM^-5 (Zymo Research).

DNA inputs of 10 ng were used with NEBNexta Ultra II FS with 6-8 cycles of PCR after 15 minutes of fragmentation and sequenced on a NextSeq 1000 P2 300 cycle cartridge for 150 bp PE reads. RNA libraries were prepared with 100 ng with Illumina Stranded Total RNA with Ribo-Zero Plus with 15 cycles of PCR on a P2 200 cycle kit for 100 bp PE reads. Initial inoculum samples were first prepared with Ribo-Zero Plus depletion to determine depletion efficiency and design custom depletion probes for the experimental dataset. The experimental dataset had 1 µL of 50 pmol supplemental probes (IDT) added at the rRNA depletion step (15 µL total volume). Best efforts were attempted in sequencing biological triplicates; however, not all samples extracted produced libraries of sufficient yield and quality for sequencing. Details on RNA probe design to carry out ribosomal RNA depletion are provided in **Supporting Methods**.

### Bioinformatic analyses

#### PacBio MAGs

Assemblies were performed for each sample’s extraction read set as well as all reads combined using hifiasm_meta v. 0.3-r063.2 (38) and metaMDBG v. 0.3 (39) and subsequently run through the pb-metagenomics HiFi-MAG-Pipeline v. 2.0.2 for binning with completeness aware binning set to 90% (40). Metagenome assembled genomes (MAGs) were all pooled for each sample and assembly type to be dereplicated using dRep v. 3.5.0 (41) with secondary average nucleotide identity (s_ANI) cluster set to 98%.

#### Short read MAGs

Sequencing reads from the experimental set up were assembled and binned using the nf-core/MAG 2.5.4 pipeline (https://nf-co.re/mag/2.5.4/). Binned MAGs were then assessed separately via checkM2 v1.0.1 and filtered to completeness cutoffs of 70% and contamination of 10%. Binned MAGs were combined with dereplicated PacBio MAGs and dereplicated again at 98% ANI for a combined set of 98 MAGs. Final classification was completed with GTDB-TK v2.40 using the R220 database. Sylph v. 0.61 (42) was used to classify the reads according to the pooled MAG set used as a reference database across the experimental treatments for comparing relative abundances.

#### Metabolic modeling

MAG annotation and metabolic summaries were produced using the Distilled Refined Annotation of Metabolism (DRAM) tool v 1.5.0 (https://github.com/WrightonLabCSU/DRAM/wiki) (43). DRAM outputs three main files that can be used in downstream analyses including a gene annotation file, a metabolic summary contained in MS Excel, and a tab-delimited file summarizing the percent completion and presence/absence data for essential metabolic pathways. We used custom scripts in R v. 4.3.2 developed using the ‘tidyverse’ package to connect the gene presence/absence data for the nitrogen metabolic module from DRAM to the relative abundance data and taxonomy data (see **Data Availability** statement for access to data analyses workflows).

#### Metatranscriptomics analysis

SqueezeMeta v. 1.6.3 (44) processed fastp v. 0.23.4 (45) cleaned DNA and RNA reads from successful libraries prepared from enrichment culture samples using a co-assembly mode on DNA reads with MEGAHIT (46, 47) and a minimum contig length of 500. KEGG annotations were queried using SQMTools v. 1.6.3 (44). Outputs from SqueezeMeta were subsequently processed in R v 4.3.2 to generate DNA and RNA read count figures and conduct data manipulations to identify unique KEGG IDs tied to the top 25 hits for read counts associated with the amino acid modules in KEGG. SqueezeMeta v. 1.7.0b8 was used for RNA read mapping back to the 98 MAGs analyzed in this work.

## Data Availability

Sequencing reads are deposited under BioProject PRJNA1165788. Metagenomic assemblies and analysis outputs are available at **10.5281/zenodo.14861211**. All R code, supplementary data files, and raw data files with reference to the figures they served as input for in this work have also been provided at **10.5281/zenodo.14861211** and on a dedicated GitHub repository (https://github.com/carleton-envbiotech/Consortium_meta_omics).

## Acknowledgments

We are sincerely grateful to our industry partner for providing us with the consortium in development that was integral to this work. This research was funded by the Chemicals Management Plan Research fund at Environment and Climate Change Canada (ECCC), Carleton University, and an NSERC Discovery Grant (2022-04891) awarded to Daniel S. Grégoire. Derek D.N. Smith also acknowledges related support for genomics research through the Genomic Research and Development Initiative on Antimicrobial Resistance 2 One Health (GRDI-AMR2 One Health) project and ECCC’s GRDI Strategic Technology Applications of Genomics in the Environment (STAGE).

